# Efficacy of oseltamivir treatment in influenza virus infected obese mice

**DOI:** 10.1101/2021.04.28.441903

**Authors:** Rebekah Honce, Jeremy Jones, Brandi Livingston, Leonardo D. Estrada, Lindsey Wang, William Caulfield, Burgess Freeman, Elena Govorkova, Stacey Schultz-Cherry

## Abstract

Obesity has been epidemiologically and empirically linked with more severe disease upon influenza infection. To ameliorate severe disease, treatment with antivirals, such as the neuraminidase inhibitor oseltamivir, are suggested to begin within days of infection, especially in hosts at higher risk for poor outcomes. However, this treatment is often poorly effective and can generate resistance variants within the treated host. Here, we hypothesized that oseltamivir treatment would not be effective in genetically obese mice and would generate a more diverse and drug resistant viral population. We demonstrated that oseltamivir treatment does not improve viral clearance in obese mice. While no traditional variants associated with oseltamivir resistance emerged, we did note that drug treatment failed to quench the viral population and did lead to phenotypic drug resistance *in vitro.* Mechanistically, we demonstrate the blunted interferon response in obese hosts may be contributing to treatment failure, as type I interferon receptor deficient mice also fail to clear influenza virus infection upon oseltamivir administration. Together, these studies suggest that the unique pathogenesis and immune responses in obese mice could have implications for pharmaceutical interventions and the within-host dynamics of the influenza virus population.

**IMPORTANCE:** Influenza virus infections, while typically resolving within days to weeks, can turn critical especially in high-risk populations. Prompt antiviral administration is crucial to mitigating these severe sequalae, yet concerns remain if antiviral treatment is effective in hosts with obesity. Here, we show that oseltamivir does not improve viral clearance in genetically obese or type I IFN receptor-deficient mice and increases the genetic entropy of the within-host viral population. This suggests a blunted immune response may impair oseltamivir efficacy and render a host more susceptible to severe disease. This study furthers our understanding of oseltamivir treatment dynamics both systemically and in the lungs of obese mice, as well as the consequences of oseltamivir treatment for the within-host emergence of drug-resistant variants.

## INTRODUCTION

Influenza viruses are a seasonal and pandemic threat to human and animal health worldwide (1). Influenza A and B viruses commonly cause seasonal outbreaks in humans, with control measures needed to mitigate its health and economic effects (2, 3). Control of influenza infection is accomplished through vaccination strategies aimed to prevent disease and antiviral courses designed to mitigate symptoms, severity, and spread upon infection (4). Several antivirals have entered clinical use, including adamantanes targeted to the M2 ion channel, neuraminidase inhibitors (NAIs) that block viral release, and endonuclease inhibitors that stall viral replication (5).

However, these antiviral treatment strategies are not always effective. First, resistance to each class of influenza antivirals has emerged, with the adamantanes no longer clinically effective due to widespread resistance (6, 7). While contemporary influenza viruses are largely susceptible to oseltamivir, many NAI resistant variants have been characterized (8–11). Second, poor host responses and delayed treatment with antivirals can impede their efficacy. Host characteristics can promote the emergence of resistant variants, as influenza infection of immunocompromised hosts results in higher rates of antiviral resistant variants (12–17).

The obesity epidemic highlights these dual concerns. Rates of worldwide obesity have nearly tripled in the past three decades (18). Obesity results in a chronic state of immunosuppression that impairs the antiviral response to infection, including the type I interferon response (19–22). Epidemiological data suggests antivirals such as oseltamivir are protective in hosts with obesity, yet empirical studies show obese mice require 10-fold higher doses to achieve complete protection (23, 24). We have previously shown that the obesogenic state impacts viral population dynamics by increasing viral diversity and promoting a more virulent influenza population (21, 25). Thus, we questioned if oseltamivir treatment of obese mice would accelerate viral clearance or lead to the acquisition of antiviral resistant variants. In these studies, we treated obese mice with the NAI oseltamivir (Tamiflu), and monitored viral clearance over 7 days. Oseltamivir treatment improved viral clearance in wild-type mice; however, no such improvement was detected in treated, leptin-deficient obese mice. Additionally, we report that this subpar protection promotes the emergence of influenza viral variants with increased neuraminidase activity and greater oseltamivir resistance. Together, these findings suggest study is warranted on how host characteristics can influence pharmaceutical efficacy.

## RESULTS

### Oseltamivir treatment does not reduce viral titers or improve viral clearance in influenza-infected obese mice

We have previously determined that higher doses of oseltamivir are needed to improve survival in obese mouse models (23, 24); however, why the standard dose treatment fails in obese mice was unknown. To answer, we asked if oseltamivir treatment could reduce viral titers or improve viral clearance in wild-type and obese mice. Beginning 12 hours pre-infection lean and obese animals were orally gavaged with 10 mg/kg oseltamivir or PBS vehicle control every 12 hours for 5 days post-infection (p.i.) (23). Mice were infected with A/California/04/2009 (CA/09) H1N1 influenza virus and lungs excised at 0.5-, 1-, 3-, 5- and 7-days p.i. (Figure 1A). No significant differences in viral titers were detected (Figure 1B), but area under the curve (AUC) analysis suggests significantly accelerated viral clearance upon oseltamivir treatment for wild-type mice compared to all other experimental groups (vs wild-type + PBS *p*=0.012; vs obese + PBS *p*=0.0007; vs obese + oseltamivir *p*=0.0014; Figure 1C). No trend in accelerated viral clearance was observed for oseltamivir-treated obese mice compared to untreated obese mice in AUC analysis. The prolonged viral replication coupled with oseltamivir pressure could result in the selection of potentially resistant variants.

**Figure 1.**
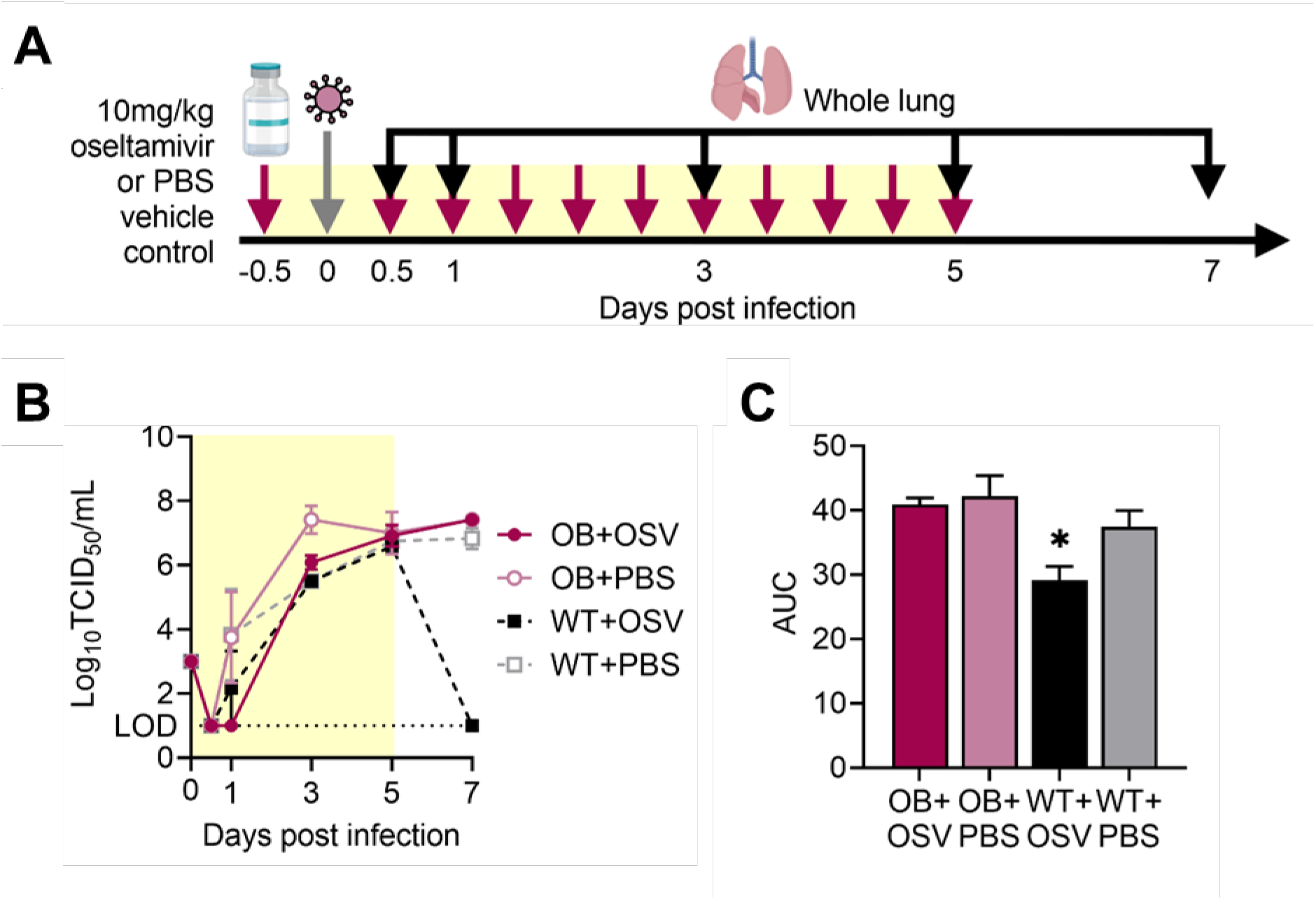
Oseltamivir treatment improves viral clearance in wild-type, but not obese, mice. (A) Male WT and OB mice were treated with 10 mg/kg oseltamivir or PBS vehicle control every 12 hours starting 12 hours pre-infection with CA/09 virus. (B) Oseltamivir treatment has no impacts on overall viral load but increased viral clearance is observed in WT mice treated with oseltamivir. (C) Area-under-the-curve analysis for viral titers in (B). Data represented as means ± SEM. Data in (B,C) represented as means ± SEM and analyzed via (B) repeated measures one-way ANOVA with Tukey’s multiple comparisons test and in (C) with ordinary one-way ANOVA with Tukey’s multiple comparisons test with α=0.05. Yellow shading indicates active treatment. OSV=oseltamivir phosphate.

### Obese-derived viruses are more resistant to oseltamivir treatment compared to those obtained from lean hosts

While emergence of antiviral resistant markers is relatively rare, we hypothesized that this delayed viral clearance could generate resistance to the drug in immunocompromised obese hosts (17, 26). To test this, we compared the neuraminidase (NA) activity of lean and obese-derived viruses at different days p.i. by MUNANA. Wild-type derived viruses have low NA activity regardless of days post-infection in the presence of oseltamivir. In contrast, the virus isolated from obese host has high NA activity by day 5 p.i. that remains high throughout the course of infection. While the presence of oseltamivir initially suppresses NA activity in obese mice, without impacting overall viral titers (Figure 1B). by day 7 p.i., viruses have emerged that have NA activity in the presence of oseltamivir. Initially, viruses derived from all four experimental groups showed similar levels of neuraminidase activity as measured by fluorescence (Figure 2A). By day 5 p.i., virus derived from untreated obese-derived virus showed greater relative fluorescence than other groups. After the removal of oseltamivir treatment, oseltamivir-treated obese mice displayed a rebound in neuraminidase activity (Figure 2A). This suggests there may be a selection of higher-NA activity viral variants within obese, oseltamivir-treated mice.

**Figure 2.**
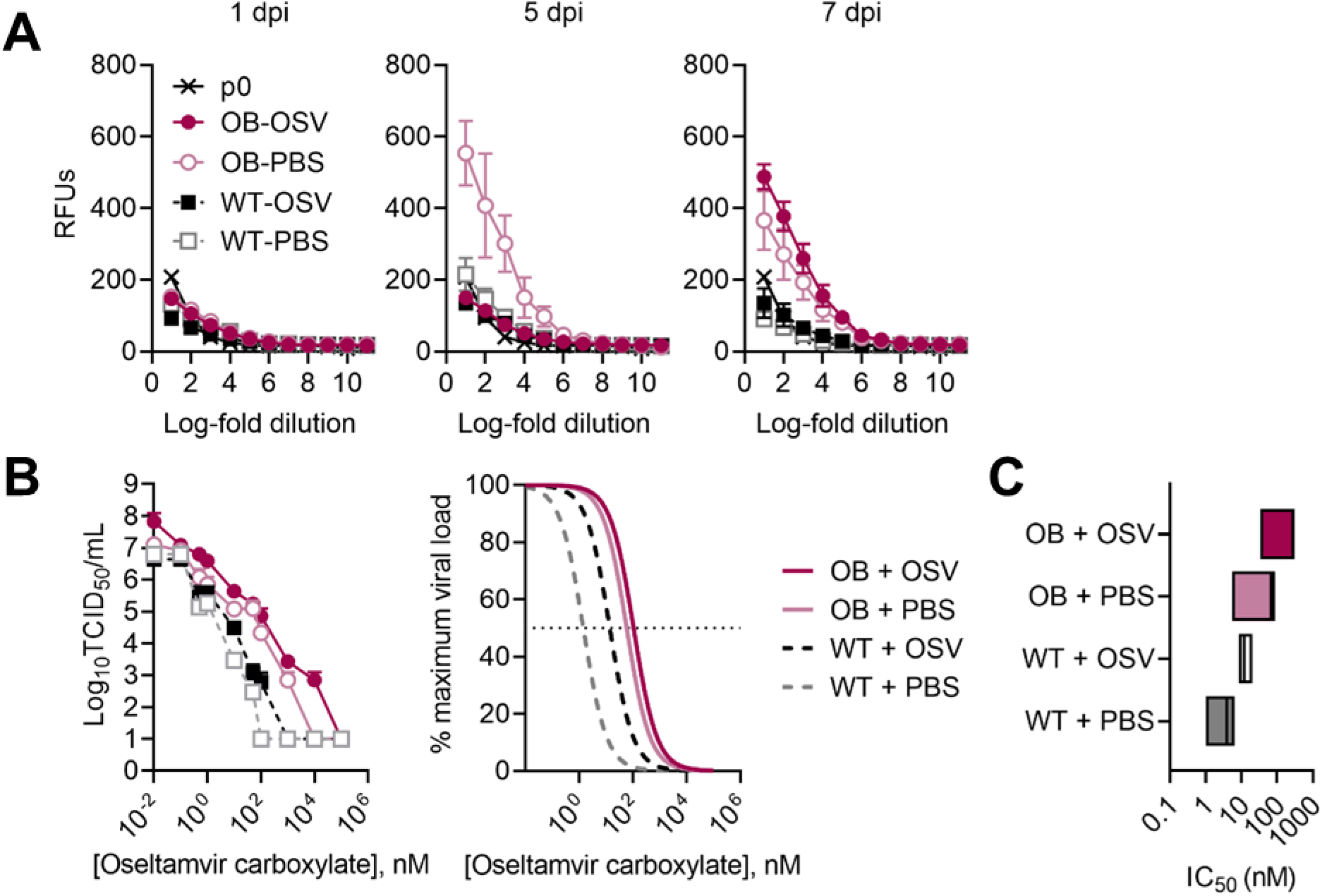
Obese-derived viruses are more resistant to oseltamivir carboxylate. (A) Relative neuraminidase activity is higher in obese-derived viruses. (B-C) Indicated viruses were titrated in the presence of increasing concentrations of oseltamivir carboxylate, with (B) viral titers determined through TCID_50_ and average non-linear curve fits of viral loads compared to maximum at no treatment, and (C) inhibitory concentrations of oseltamivir carboxylate needed to reduce viral load by 50% compared to no treatment. Data presented as (A,B) means ± SEM and analyzed in (B) through area-under the curve analysis via one-way ANOVA with Tukey’s multiple comparisons test and non-linear fits with a non-linear fit curve to determine IC_50_s reported in (C). OSV=oseltamivir phosphate.

To quantitate the difference in potential oseltamivir resistance between wild-type and obese derived-viruses, the indicated viruses were titrated on MDCK cells in the presence of DMSO control or increasing concentrations of oseltamivir carboxylate as indicated. Viruses derived from obese, treated mice trended towards higher titers at all experimental conditions compared to both treated and untreated wild-type-derived viruses (Figure 2B). AUC analysis showed significantly reduced viral titers across all experimental conditions in treated wild-type (*p*=0.0316) and all untreated (wild-type + PBS, *p*=0.0173; obese + PBS *p*=0.0425) mice compared to treated obese mice (obese + oseltamivir). The half-maximal inhibitory concentration (IC_50_) for oseltamivir ranges from 0.8 nM to greater than 35 μM (6, 12). Within our experimental conditions, (IC_50_) values for all obese-derived viruses were shifted higher compared to wild-type derived viruses, with oseltamivir -treatment further shifting resistance in both groups (Figure 2C). Mean IC_50_ values are as follows: obese + oseltamivir, 206.4 ± 86.3 nM; obese + PBS, 52.8 ± 24.4 nM, wild-type + oseltamivir, 13.7 ± 3.5 nM, wild-type + PBS 3.8 ± 1.6 nM. In total, obese-derived viruses show greater phenotypic resistance to oseltamivir carboxylate treatment *in vitro* compared to lean-derived viruses.

### Obese-derived viruses do not have genetic changes commonly associated with antiviral resistance

The influenza virus hemagglutinin (HA) and NA segments are genetically plastic allowing for accrual of potentially resistant single nucleotide variants (SNVs). The HA mutations G155E and D222G as well as the NA mutation H275Y are implicated in reducing oseltamivir efficacy, with several others also suggested as modulators of resistance (27, 28). To determine if these mutations were present in the obese-derived viruses, NGS was performed and we quantified the overall number of single nucleotide variants SNVs, entropy, and if any classically NAI-resistant mutants emerged. No consensus changes associated with oseltamivir resistance were detected.

Viruses derived from obese mice treated with oseltamivir had significantly increased numbers of unique SNVs compared to viruses derived from wild-type mice treated with oseltamivir (*p*=0.0053; Figure 3A). This translated to increased overall genetic diversity in viruses derived from oseltamivir-treated obese mice. Measures of Shannon’s entropy (H) is reduced in wild-type treated (*p*=0.0035) and wild-type untreated (*p*=0.0357) compared to oseltamivir treated obese mice (Figure 3B). Oseltamivir ablated overall viral diversity in lean mice. There was a non-significant trend to reduced numbers of SNVs and total Shannon’s entropy in treated lean mice, compared to no difference in treated obese mice (Figure 3A, B).

**Figure 3.**
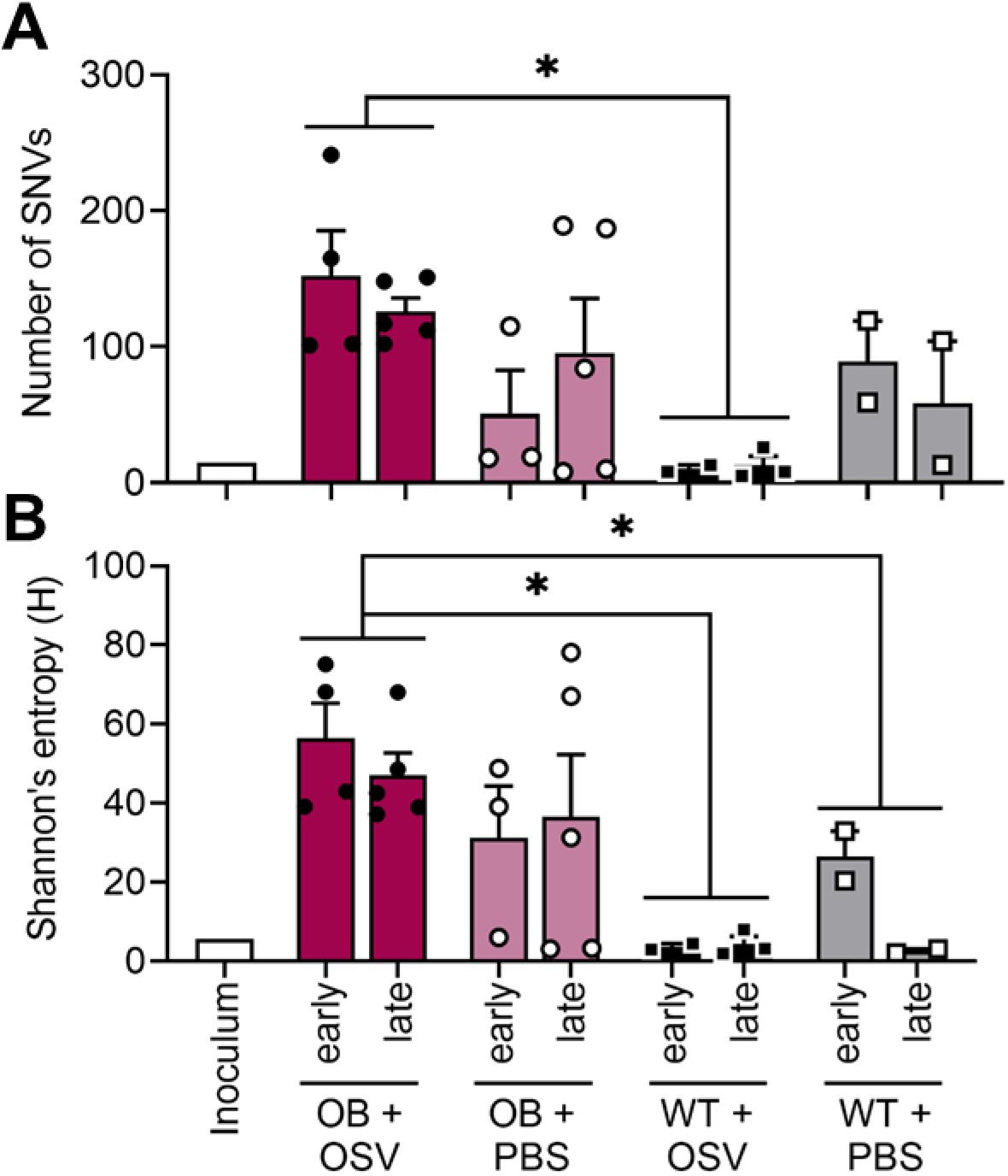
Oseltamivir treatment ablates viral diversity in wild-type, but not obese, mice. (A-B) Viral RNA was extracted from lungs of mice treated with oseltamivir as described in Figure 2A. Total unique SNVs are significantly higher in oseltamivir treated OB mice compared to treated WT mice (*p*=0.0053). (B) Oseltamivir treated OB mice harbor a more diverse viral population compared to WT Oseltamivir treated (p=0.0035) and PBS-control treated (p=0.0357) mice. Statistical comparisons made via two-way ANOVA with Tukey’s multiple comparisons test. Significance was set at α=0.05. Data represented as means ± SEM. OSV=oseltamivir phosphate.

More SNVs were detected in the HA segment of viruses derived from obese compared to lean oseltamivir treated mice (9 and 4, respectively) translating to higher average entropy of 3.27 in obese mice compared to 1.24 in wild-type mice. Oseltamivir treatment decreased HA entropy in treated wild-type mice (from 1.98 to 1.24) but not in obese mice, where it increased from 2.30 to 3.27 (Table 1). We detected fewer SNVs in NA. Oseltamivir treatment resulted in 1 NA variant in obese and 3 NA variants in lean, compared to 2 and 1 in obese and lean PBS control mice, respectively (Table 1). While we noted higher genetic diversity and phenotypic resistance in both oseltamivir-treated and untreated mice, no classical NAI-resistant mutants emerged in the viral populations. However, there is no exhaustive list of mutations than may render oseltamivir treatment less effective. Combined with the increased rate of replication and viral population diversification in obese mice, this may be the ideal context for emergence of novel resistance markers.

**Table 1.**
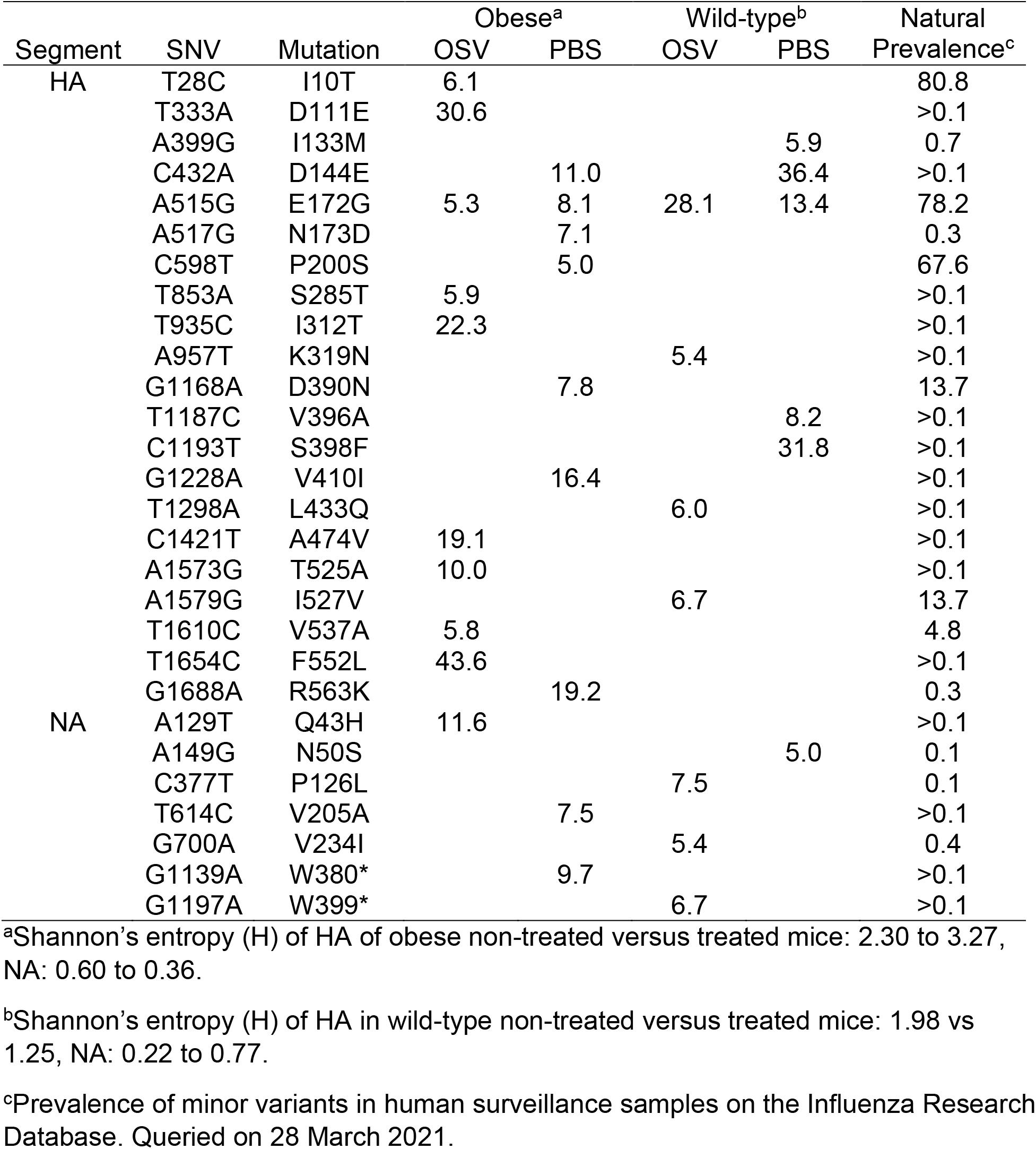
Percentage of detected minor variants in obese and wild-type mice treated with oseltamivir.

### Similar oseltamivir clearance but reduced maximal concentrations in obese compared to lean mice

Appropriate dosage and concentration at the site of infection is crucial for antiviral efficacy; inappropriate levels could lead to severe complications due to treatment failure or the quick emergence of resistance phenotypes. To determine if obesity impacts the pharmacokinetic dynamics of oseltamivir and the metabolite oseltamivir carboxylate, we dosed male, 10-week-old wild-type or obese mice with a single oral gavage of oseltamivir at 10 mg/kg in 100 uL of PBS. Whole lungs and plasma were collected at 0.5-, 1-, 4-, 8-, and 16-hours post treatment and immediately processed for pharmacokinetic (PK) analysis (Figure 4A). There were no practical or significant differences in oseltamivir or oseltamivir carboxylate clearance in plasma and lung between wild-type and obese mice (Table 2). We observed a similar half-life of oseltamivir (2.12 versus 2.28 hours; Figure 4B) and its metabolite oseltamivir carboxylate (2.30 versus 3.3 hours; Figure 4C) in the lungs of obese mice compared to wild-type mice. Analysis of plasma revealed similar findings, with no difference in the elimination half-life of oseltamivir (1.36 versus 1.98 hours, respectively; Figure 4E) or oseltamivir carboxylate (1.87 versus 2.75 hours, respectively; Figure 4F) in lean versus obese mice. However, the maximum concentration of oseltamivir was significantly increased in both plasma and lung tissue. In lungs, maximal oseltamivir concentration rose from an average of 600 ug/L in obese mice lungs to 1500 ug/L in the lungs of wild-type mice (*p*=0.036; Figure 4D). Plasma again showed similar trends (*p*=0.063; Figure 4G). Overall, these findings suggest that clearance of oseltamivir is equal between obese and lean mice and most likely has little impact on the observed failure of viral clearance.

**Figure 4.**
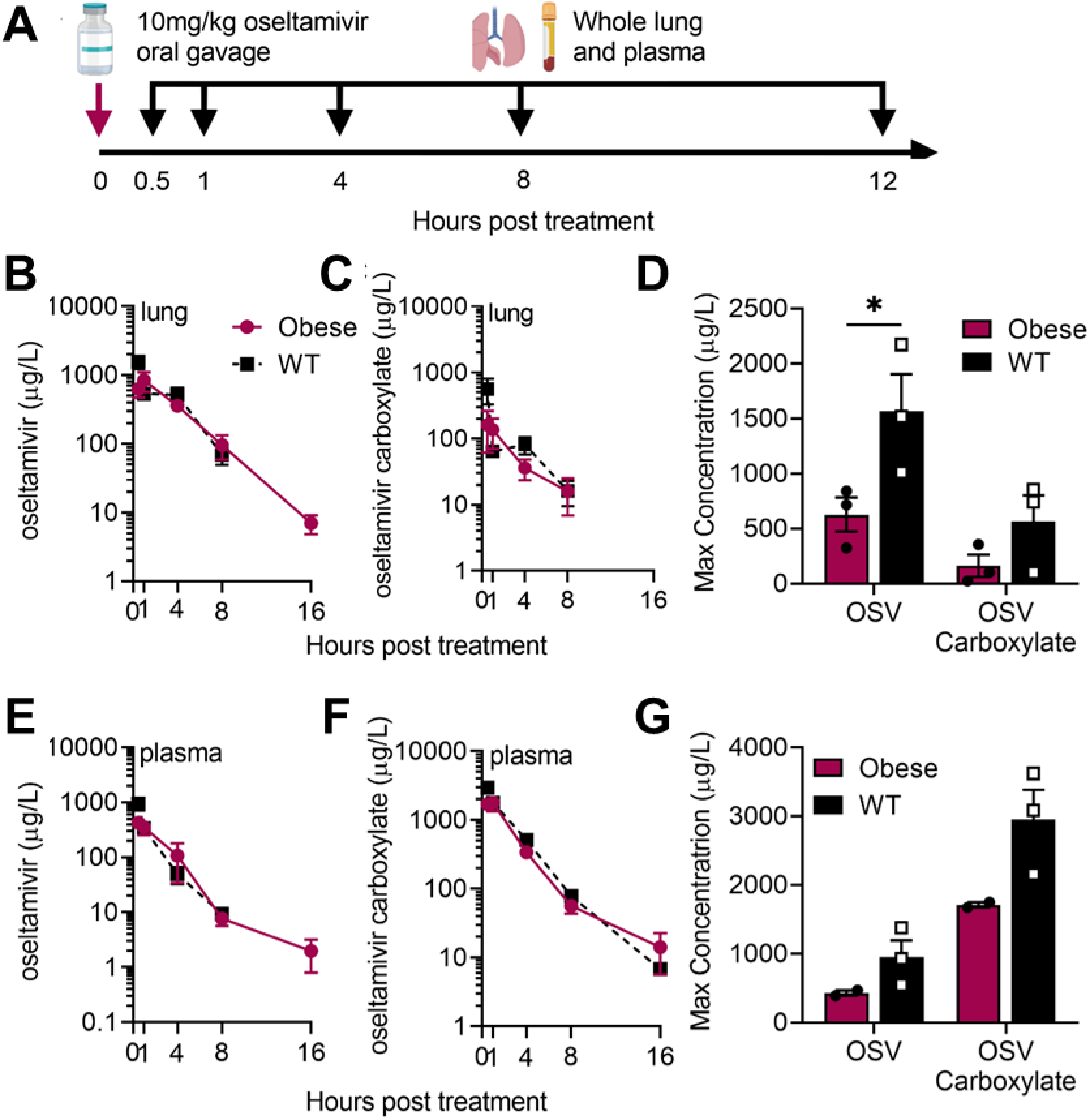
Similar oseltamivir pharmacokinetic parameters in lean and obese mice. (A) Naïve male, WT or OB mice were dosed once with 10 mg/kg of oseltamivir. Plasma and lungs were collected at indicated time points for bioanalysis of (B) oseltamivir and (C) oseltamivir carboxylate in lungs. (D) Significantly increased maximum concentration of oseltamivir (*p*=0.0364) and trends towards increased oseltamivir carboxylate levels was observed in the lungs of WT compared to OB mice. (E-F) After one dose of oseltamivir, bioanalysis of (E) oseltamivir and (F) oseltamivir carboxylate in plasma of WT and OB mice. (D) Maximum concentration of oseltamivir and oseltamivir carboxylate in plasma of WT and OB mice. Non-significant trend towards increased concentration of oseltamivir and oseltamivir carboxylate in WT compared to OB plasma. Data analyzed in (B, C, E, F) with ordinary two-way ANOVA and in (D, G) with two-way ANOVA and Sidak’s post hoc with α=0.05. Data represented as means ± SEM.

**Table 2.**
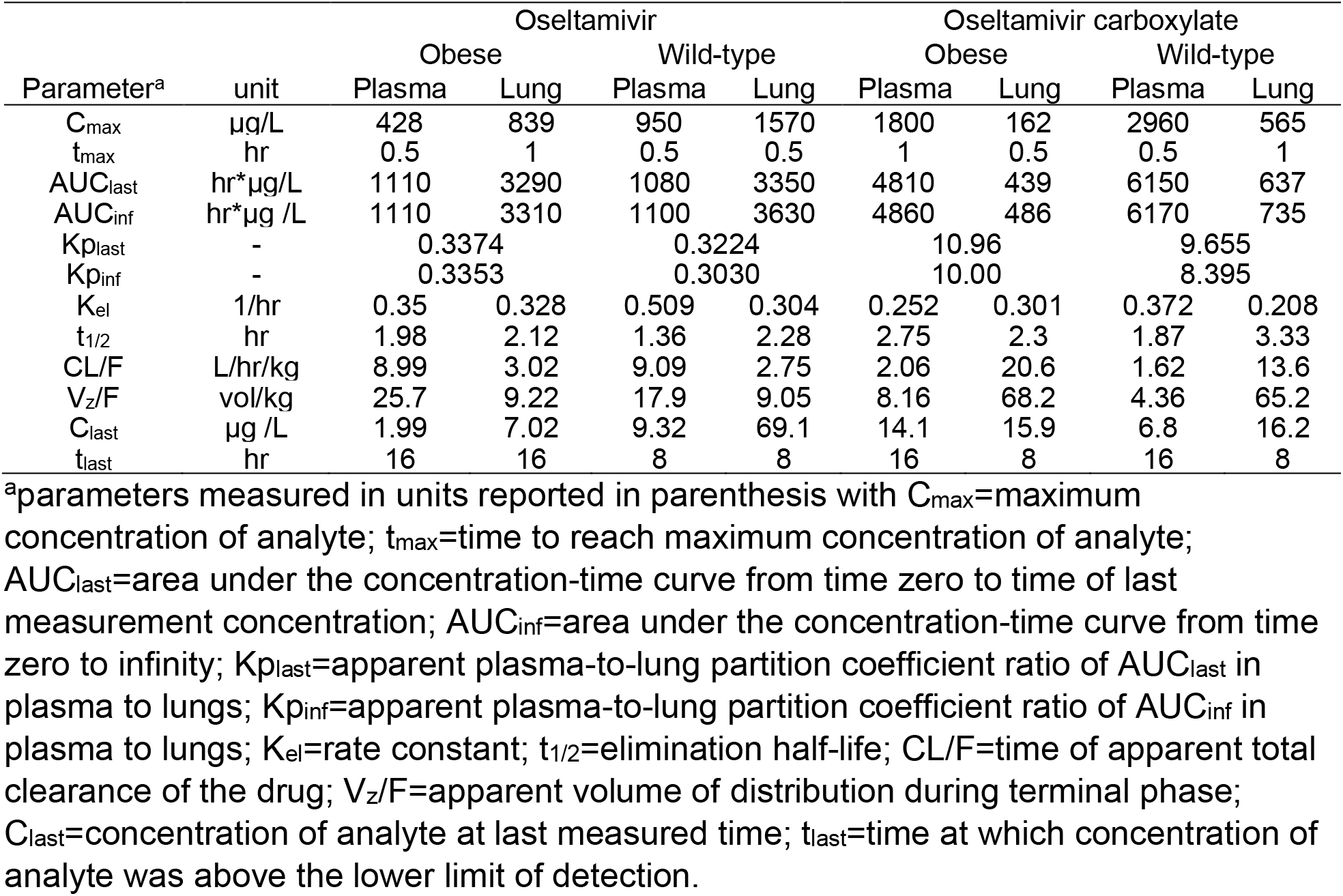
Pharmacokinetic parameters of oseltamivir and oseltamivir carboxylate in plasma and lungs of obese and wild-type mice.

### Interferon is important for oseltamivir control of influenza virus infection

Obesity is associated with reduced type I (IFN) responses. We further demonstrated that robust IFN responses restrict genetic diversity. To test the hypothesis that a reduced IFN response can decrease the effectiveness of oseltamivir due to increased phenotypically resistant variants, male and female IFNAR^−/−^ or WT mice were treated with 10 mg/kg oseltamivir or vehicle control (PBS) and inoculated intranasally with CA/09 H1N1 virus or PBS as in Figure 1A. Like obese mice (Figure 1B-C), oseltamivir treatment showed little effectiveness in reducing viral load in IFNAR-null mice (Figure 5A). Only, wild-type mice treated with oseltamivir had reduced viral clearance. IFNAR^−/−^ mice, with and without oseltamivir treatment, had detectable pulmonary viral loads at both days 3 and 7 p.i. This suggests that IFN signaling is required for robust oseltamivir antiviral activity and improved viral clearance in mice, even when equal dosage and pKa dynamics point towards treatment efficacy.

**Figure 5.**
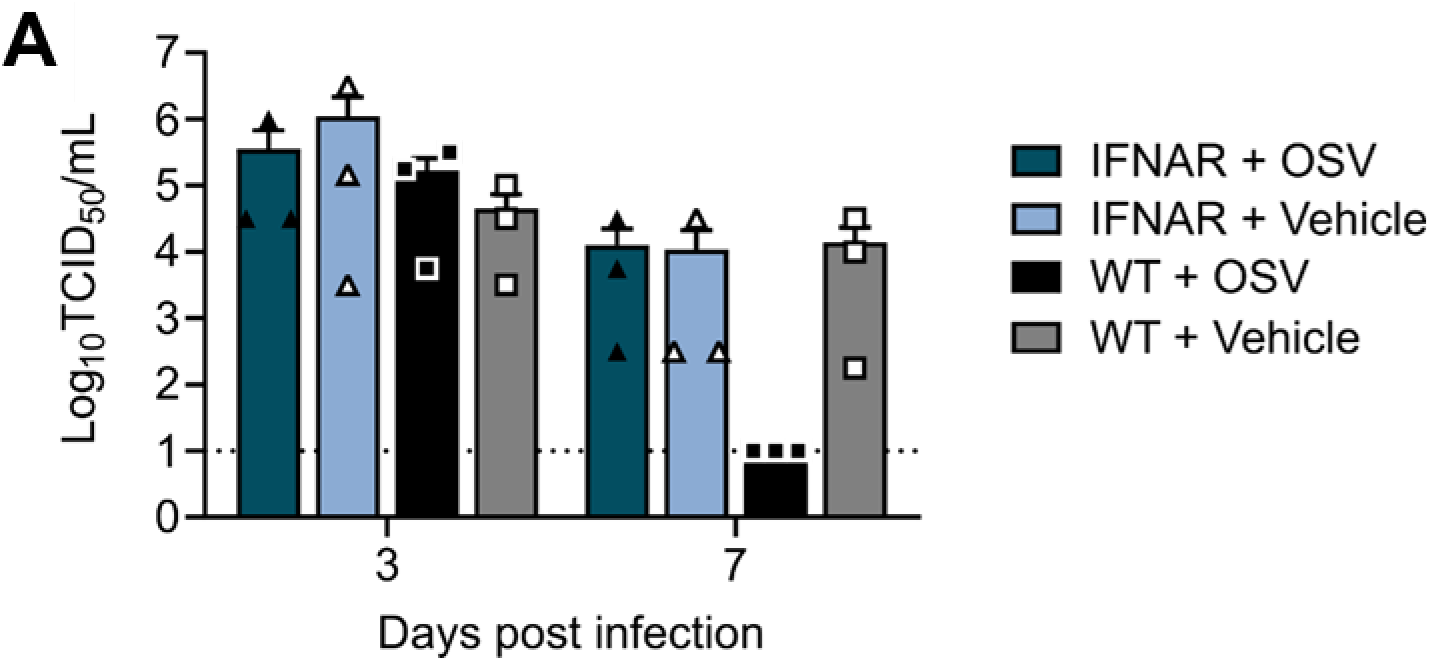
Interferon-deficient animals are not responsive to oseltamivir treatment. WT and IFNAR^−/−^ mice treated with OSV or vehicle (PBS) and inoculated with CA/09 or mock (PBS) were monitored for 7 days. (A) Viral load in lungs at days 3 and 7 p.i. with CA/09 virus. Data in represented as means ± SEM and analyzed via ordinary one-way ANOVA with Tukey’s multiple comparisons with α=0.05. OSV=oseltamivir phosphate.

## DISCUSSION

In these studies, we demonstrate that phenotypic resistance to oseltamivir emerged by day 5 p.i. in obese mice during treatment. We hypothesize that the delayed type I IFN response, in conjunction with reduced maximum concentration of oseltamivir in the pulmonary environment, may result in poor control of viral replication allowing the emergence of resistant viral variants. Population-based pharmacokinetic studies have found that oseltamivir clearance is accelerated in obese hosts, but not at a biologically significant rate spurring no need for dosage based on weight (24, 27–29). However, these studies have relied on plasma concentrations and not the concentration at the respiratory epithelium. No studies to our knowledge have either empirically tested antiviral efficacy in humans with obesity or the local concentration of oseltamivir at the site of infection.

Emergence of dominant antiviral resistant mutations often begin as minor variants in the within-host population, especially in infected, immunocompromised hosts (30, 31). Compounding these risks, oseltamivir-resistant variants in already immunocompromised hosts, such as those with obesity, may complicate an already high-risk medical presentation (32). Weakened immune pressures, including blunted innate and adaptive immunity, coupled with extended shed and potentially higher viral replication may increase the likelihood of adaptive mutations emerging in obese hosts (33). While paradoxically, oseltamivir treatment is not required for the emergence of oseltamivir-resistant variants, adding on this selection pressure in an already compromised situation may prove a perfect storm for viral adaptation. To remedy this, early and appropriate antiviral responses are crucial to control of viral replication, spread and pathology. This is mirrored by the window in which oseltamivir is effective in preventing the onset of severe disease (23). While those with obesity are more at-risk for severe sequalae following influenza infection, prompt antiviral administration may ameliorate these risks and improve overall outcomes (34). Treatment delayed as little as 2 days after symptoms onset is ineffective in most hosts, mirroring the sensitive timing for robust action of endogenous IFN responses (4, 35).

We have previously shown that IFN is crucial for controlling the emergence of a diverse viral population that may harbor virulent genotypes (21). Type I IFN and oseltamivir may also show synergistic benefit in treating seasonal IAV infection, and oseltamivir has been shown to modulate immune responses (36–38). The blunted pulmonary IFN responses coupled with reductions in maximal oseltamivir concentrations in obese hosts may increase the likelihood for antiviral resistant variants to emerge leading to the observed phenotypic resistance (23, 39, 40). Identifying the molecular mechanisms behind the blunted IFN responses will be crucial to untangling the multiple impacts the obesity epidemic has on public health.

## AKNOWLEDGEMENTS

This manuscript has benefitted from helpful discussion with Drs. Nicholas Wohlgemuth and Victoria A. Meliopoulos. This work was supported by the National Institute of Allergy and Infectious Diseases under HHS contract HHSN27220140006C for the St. Jude Center of Excellence for Influenza Research and Surveillance funding from the National Institutes of Health, NIH grant R01AI140766, and ALSAC.

## METHODS

### Viruses and titer determination

Eight- to 12-week old mice were inoculated with indicated doses of A/California/04/2009 (H1N1) virus and viral titer determined through tissue-culture infectious dose-50 (TCID_50_) assays as previously reported (23).

### Animal husbandry

Eight to 12-week old wild-type C57Bl/6 male (wild-type) (JAX:000664) and B6.C-Lep^*ob/ob*^/J genetically obese (obese) (JAX:000632) male mice were obtained from Jackson Laboratory. IFNAR KO mice were obtained from Dr. Laura Knoll (University of Wisconsin) and bred in house. Knockouts were confirmed by PCR using primer sets (IFNAR^−/−^) reported on the Jackson Laboratories website. All animals were housed under standard conditions with food and water provided *ad libitum.* All procedures were approved by the St. Jude Children’s Research Hospital Institutional Animal Care and Use Committee and followed the Guide for the Care and Use of Laboratory Animals.

### In vivo pharmacokinetics (PK)

Plasma and lung tissue pharmacokinetic (PK) profiles of prodrug oseltamivir and active metabolite oseltamivir carboxylate was evaluated in male C57BL6 and obese mice (Jackson Labs), approximately 12 weeks in age. Oseltamivir phosphate was dissolved in PBS at 5 mg/mL for a 10 mL/kg gavage, yielding a 10 mg/kg dose. Terminal blood samples were obtained at various times up to 16 hours post-dose and immediately processed to plasma. A 200 μL aliquot of plasma was then quickly pipetted from each sample and transferred into a separate 2 mL microcentrifuge tube containing 800 μL of ice cold 15 ng/mL oseltamivir-d3 carboxylate (Toronto Research Chemicals, Lot 14-SBK-47-3, Purity 97%) in methanol, capped, vortexed for 30 seconds and then centrifuged to pellet the precipitated protein. An 800 to 950 μL aliquot of the supernatant was then pipetted into an empty pre-labeled microcentrifuge tube, capped and stored at −80 °C until analysis. Following terminal bleeds, animals were perfused with PBS to flush blood from the vasculature. Lungs were then extracted, rinsed with PBS as necessary, and snap frozen in liquid nitrogen. Lungs were then placed in appropriately labeled microcentrifuge tubes in a cooler on dry ice and transferred to −80°C for storage.

### Bioanalysis

Frozen lung samples were weighed in tared 2 mL Lysing Matrix A (MP Biomedical, Santa Ana, CA) tubes and diluted with a 5:1 volume of LCMS grade methanol. The lungs were then homogenized with a FastPrep-24 system (MP Biomedicals, Santa Ana, CA). The homogenization consisted of three 6.0 M/S vibratory cycles of 1 min each on the FastPrep-24 system. To prevent over-heating due to friction, samples were placed on wet ice for 5 min between each cycle. The homogenates were then centrifuged at 14,000 x g and stored at −80 °C until analysis. Plasma and lung homogenate samples were analyzed for osetalmivir (oseltamivir phosphate, Cayman Chemical Co., Batch 0458276-29, purity 98%) and oseltamivir carboxylate (Toronto Research Chemicals, Lot 3-SKC-52-1, Purity 98%) with a qualified LC MS/MS assay. Calibration and quality control stock solutions were prepared in DMSO and serially diluted in DMSO to prepare calibration/QC spiking solutions. For the analysis of plasma samples, these spiking solutions were then used to prepare calibration and QC samples in methanol. Methanol calibration and QC samples, 200 μL each, were then pipetted into glass HPLC vials and evaporated on a CentriVap Console (Labconco) evaporator (40 minutes at 60 °C, then 30 minutes at 70 °C). The residue was reconstituted in 800 μL of ice cold 15 ng/mL osetalmivir-d3 carboxylate in methanol as an internal standard. Blank male C57Bl/6 plasma (200 μL) was then pipetted into each vial, immediately vortexed for 1 minute and centrifuged to pellet the protein.

Since the lungs were simultaneously homogenized and protein precipitated in methanol, the calibration and QC samples were prepared by spiking into blank male C57Bl/6 lung homogenate. Aliquots (25 μL) of the standards, QC solutions and samples were then pipetted into the IS spiking solution and vortexed to mix. A 2 μL aliquot of the extracted supernatant was injected onto a Shimadzu LC-20ADXR high performance liquid chromatography system via a LEAP CTC PAL autosampler. The LC separation was performed using a Phenomenex Kinetex Polar C18 (2.6 μm, 50 mm x 2.1 mm) column maintained at 50 °C with gradient elution at a flow rate of 0.50 mL/min. The binary mobile phase consisted of water-formic acid (100:0.1 v/v) in reservoir A and acetonitrile-formic acid (100: 0.1 v/v) in reservoir B. The initial mobile phase consisted of 5% B with a linear increase to 55% B in 3 min. The column was then rinsed for 2 min at 100% B and then equilibrated at the initial conditions for 2 min for a total run time of 7 min. Under these conditions, oseltamivir carboxylate, IS and oseltamivir eluted at 1.22, 1.22 and 1.83 min, respectively.

Analyte and IS were detected with tandem mass spectrometry using a SCIEX 5500 QTRAP in the positive ESI mode and the following mass transitions were monitored: oseltamivir carboxylate 285.18 → 138.10, oseltamivir-d3 carboxylate 288.20 → 139.20 and oseltamivir 313.20 → 166.20. The method qualification and bioanalytical runs all passed acceptance criteria for non-GLP assay performance. A linear model (1/X2 weighting) fit the calibrators across the 1.00 to 500 ng/mL range, with a correlation coefficient (R) of ≥ 0.9973. The lower limit of quantitation (LLOQ), defined as a peak area signal-to-noise ratio of 5 or greater verses a matrix blank with IS, was 1.00 ng/mL. Sample dilution integrity was confirmed. The plasma intra-run precision and accuracy was ≤ 6.63% CV and 95.8% to 109%, respectively. The lung homogenate intra-run precision and accuracy was ≤ 4.48% CV and 94.9% to 111%, respectively.

### Pharmacokinetic (PK) analysis

Oseltamivir plasma Ct data were grouped by matrix and nominal time point, and the mean Ct values were subjected to noncompartmental analysis (NCA) using Phoenix WinNonlin 8.1 (Certara USA, Inc., Princeton, NJ). The extravascular model was applied, and area under the Ct curve (AUC) values were estimated using the “linear-up log-down” method. The terminal phase was defined as at least three time points at the end of the Ct profile, and the elimination rate constant (K_el_) was estimated using an unweighted log-linear regression of the terminal phase. The terminal elimination half-life (T_1/2_) was estimated as 0.693/K_el_, and the AUC from time 0 to infinity (AUC_inf_) was estimated as the AUC to the last time point (AUC_last_) + C_last_ (predicted)/K_el_. Other parameters estimated included observed maximum concentration (C_max_), time of C_max_ (T_max_), concentration at the last observed time point (C_last_), time of C_last_ (T_last_), apparent clearance (CL/F = Dose/AUC_inf_), and apparent terminal volume of distribution (Vz/F). The apparent plasma-to-lung partition coefficient (Kp_inf_) was estimated as the ratio of the AUC_inf_ in tissue to AUC_inf_ plasma, whereas Kp_last_ was similarly estimated using AUC_last_ values.

### Antiviral resistance *in vivo*

Oseltamivir was administered by oral gavage at 10 mg/kg free-base equivalencies twice daily for 5 days. At 12 hours after the initial dose, mice were lightly anesthetized with isoflurane and inoculated with 10^3^ TCID_50_ units of A/California/04/2009 (H1N1) virus in 25 μL PBS. Control mice received a PBS oral gavage at the same timepoints. At 0.5-, 1-, 3-, 5- and 7-days post infection, lungs were collected, homogenized in 1 mL PBS and stored at −80°C for downstream viral titer determination and deep sequencing. For deep sequencing statistical analysis, days 1 and 3 and days 5 and 7 are grouped as early and late points in infection, respectively, due to low copies of viral RNA at very early and very late periods in infection.

### Drug susceptibility assays

Madin-Darby canine kidney cells (MDCK cells; RRID: CVCL_0422) were maintained in minimum essential medium (MEM; Lonza) supplemented with 2 mM GlutaMAX (Gibco) and 10% fetal bovine sera (FBS; Atlanta Biologicals) and grown at 37°C under 5% CO_2_. MDCK cells were seeded in 12-well or 96-well cell culture treated plate. Upon confluency, MDCK cells in 96-well plates were inoculated in triplicate with 10-fold serial dilutions of indicated viruses in infection media (MEM, 2 mM GlutaMAX (Gibco), 1% BSA, and 1 μg/ml TPCK-treated trypsin) containing increasing concentration of oseltamivir carboxylate (0, 0.1, 0.5, 1, 10, 50, 100, 500, 1000, and 10,000 nM) or DMSO. Plates were maintained at 37°C under 5% CO_2_ for three days, at which time viral titer determined through hemagglutination of 0.5% turkey red blood cells in PBS. Infectious viral titers were calculated using the Reed-Muench method (41). The dose-response curve was fitted to determine the necessary concentration of oseltamivir to reduce viral titers by 50% (IC50). Assays were repeated three times with average IC50 reported. For modified plaque assays in 12-well plates, MDCK cells were washed twice with PBS, then inoculated at a MOI=0.01 with indicated viruses. After a 1-hour adsorption, cells were washed twice with PBS and overlaid with an 1.2% agarose in DMEM mixture containing TPCK-trypsin and increasing concentrations of oseltamivir carboxylate as indicated. At day 3 p.i., overlay was removed, and cells stained with crystal violet to visualized cytopathic effects (CPE).

### Quantifying neuraminidase activity

The relative neuraminidase activity of the oseltamivir-treated and untreated obese and wild-type-derived viruses was measured by using the fluorogenic substrate MUNANA (Sigma-Aldrich, St Louis, MO). Two-fold dilutions of day 1, 5, and 7 p.i. lung homogenates were prepared in enzyme buffer (32.5 mM of 2-(N-morpholino) ethanesulfonic acid (MES), 4 mM of calcium chloride, pH 6.5) and added in duplicate to a flat-bottom 96-well opaque black plate (Corning). Pre-warmed MUNANA substrate was added to all wells (30 μL/well) to achieve a final concentration of 100 μM. Immediately after adding the MUNANA substrate, the plate was incubated for one hour at 37°C. After incubation, the reaction was terminated by addition of 150 uL per well of stop solution (0.014M NaOH in 83% EtOH) and the fluorescence was read at excitation at 355 nm and emission at 460 nm (BioTek). Background-corrected relative fluorescence units were compared to a six-point standard curve of 4-MU.

### Deep sequencing and bioinformatics

Viral RNA was extracted from 50 μL of whole lung homogenate or NHBE cell lysates and supernatant on a Kingfisher Flex Magnetic Particle Processor (Thermo Fisher Scientific) by using the Ambion MagMAX-96 AI/ND Viral RNA Isolation kit (Applied Biosystems, cat#AM1834). RNA concentration was measured spectrophotometrically (Nanodrop). Multi-segment polymerase chain reaction (MS RT-PCR) was performed using SuperScript IV One-Step RT-PCR System with Platinum™ Taq High Fidelity DNA Polymerases (ThermoFisher, cat#12574-035) and influenza-specific universal set of primers(44) (Opti-F1 5’-GTTACGCGCCAGCAAAAGCAGG-3’, Opti-F2 5’-GTTACGCGCCAGCGAAAGCAGG-3’, Opti-R1 5’-GTTACGCGCCAGTAGAAACAAGG-3’). RNA (5 uL) was added and placed into a thermocycler paused at 55°C. The following cycling parameters were followed: 1 cycle of 55°C/2min; 1 cycle of 42°C/60min; 94°C/2min; 5 cycles of 94°C/30s, 44°C/30s, 68°C/3.5min; 26 cycles of 94°C/30s, 57°C/30s, 68°C/3.5min; 1 cycle of 68°C/10min; and then held at 4°C. 5 μL of the reaction was analyzed by 0.8% agarose gel electrophoresis to verify all genomic segments are present, and the reaction purified using the Agencourt AMPure XP Kit (Beckman Coulter) according to manufacturer’s instructions. The concentration of the purified DNA was measured spectrophotometrically prior to storage at −20°C (Nanodrop). DNA amplicons were deep sequenced using Illumina MiSeq technology performed by the St. Jude Children’s Research Hospital Hartwell Center with DNA libraries prepared using Nextera XT DNA-Seq library prep kits (Illumina, cat#FC-131-1024) with 96 dual-index bar codes and sequenced on an Illumina MiSeq personal genome sequencer. Single nucleotide variants (SNVs) relative to the reference sequence (A/California/04/2009 (H1N1)) were determined in by mapping reads using the low-variant detection method in CLC Genomics Workbench 12 (42). To determine whether the variants identified have been previously detected in human surveillance samples, we used the protein sequence variance analysis for HA and NA at the Influenza Research Database (43). Comparison of variants was made in reference to A/California/04/2009 virus, and relative variant frequencies were calculated by dividing the number of the amino acid variants by the total number of sequences queried.

### Statistical analyses and data visualization

Data were organized in Microsoft Excel and GraphPad Prism 8. Experiment schematics in Figures 1 and 5 were created using bioRender. Specifics of statistical details for each experiment can be found in the figure legends. All data are displayed as means ± standard error of the mean with stars used to denote statistical significance.

Significance was set at α=0.05.

